# Do 5’ regions of human protein-coding genes contain the blueprints for alternative splicing?

**DOI:** 10.1101/2025.11.01.686005

**Authors:** Thomas Weber, Luc Moulinier, Nicolas Scalzitti, Julie D. Thompson, Kirsley Chennen, Olivier Poch

## Abstract

To investigate alternative splicing capacity, we statistically compared the properties of human protein-coding genes with multiple transcript isoforms (MISOG) and single transcript isoforms (SISOG). Apart from global exon content, differential features are concentrated in the 5’ gene regions, with MISOG presenting complex 5’ untranslated region architecture and a distinctive flanking environment around first 5’ intron. Importantly, we found that 5’ exons are more prone to alternative splicing in MISOG. These results unravel previous observations indicating the importance of 5’ gene regions in some transcriptional processes and call for their reassessment in light of the MISOG/SISOG profiles.

## Background

Alternative splicing (AS) (1,2) in eukaryotes is a powerful process to increase a gene’s functional diversity by combining different structures and functions into distinct protein isoforms. Thanks to next generation sequencing advances (3), many human isoforms are now available, resulting in an extensive and comprehensive catalogue (4) suitable for the study of isoforms and the understanding of their importance in fundamental processes such as diseases and environmental responses (5). However, to our knowledge, there is no systematic study evaluating the complete set of Human Protein Coding Genes (HPCG) in terms of their ability to produce multiple transcript isoforms. Here, we present a robust multi-level statistical comparison of gene properties, such as the number and length of exons, untranslated regions (UTR), translated exonic regions (TER) and introns, to evaluate whether MISOG and SISOG have different profiles.

## Results and discussion

To focus on transcript isoforms of the 19,285 HPCG in the RefSeq database (4), we excluded 1,983 single-TER genes, including 1,610 SISOG and 373 MISOG with AS only in UTRs. The remaining 17,302 genes are composed of 10,995 MISOG and 6,307 SISOG, implying that MISOG are 1.7 times more frequent than SISOG. It is to note that, while 95% of genes have been estimated to undergo alternative splicing (6), only 64% of HPCG present multiple transcript isoforms curated by experts. The final gene set corresponds to 59,290 distinct transcripts (52,983 MISOG /6,307 SISOG) comprising 230,664 exons (165,585 /65,079), 213,004 TERs (150,276 /62,728) and 223,564 introns (164,826 /58,738) (Tables S1-3).

In terms of median gene lengths, MISOG (36,988 bp) are significantly longer than SISOG (20,617 bp) (Fig. 1,Table S4) and the total genomic length of MISOG (933 Mbp) is 2.9 times greater than SISOG (322 Mbp). This indicates that the larger genomic length of MISOG is not only related to their greater number (1.7 times) but also to distinct MISOG/SISOG gene architecture.

**Figure 1.**
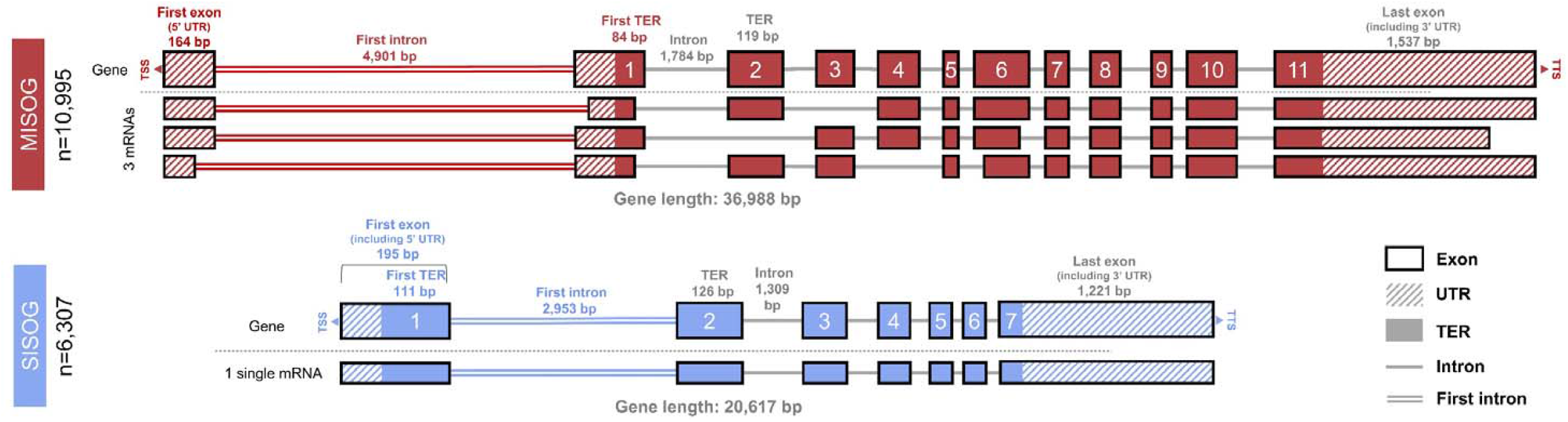
MISOG and SISOG archetypal gene architecture. Major values discussed in the manuscript are summarized graphically (all values are available in the supplementary Tables). The median lengths of each gene component (exon, TER, intron, UTR) are described according to the values observed for the MISOG (red; median number of 3 mRNAs as transcript isoforms) and SISOG (blue). The major MISOG and SISOG discriminant elements (first 5’ introns, 5’ exons and 5’ TER) are indicated by colored labels. Non-discriminant elements are indicated by grey labels.Gene lengths was calculated from TSS (Transcription Start Site) to TTS (Transcription Termination Site). TER stands for Translated Exonic Regions, the coding part for amino acids of exons.

We then performed a detailed statistical analysis of the exon, TER, UTR and intron components. The median number of exons (Fig. 1,Table S5) is significantly greater for MISOG (12 exons) than SISOG (7 exons). For SISOG, the median number of TERs is also 7, while for MISOG, the median of 11 TERs suggest additional 5’ or 3’ UTR exons. Considering median lengths (Table S6), MISOG exons (134 bp) and TERs (119 bp) are only slightly shorter than SISOG exons (136 bp) and TERs (126 bp). Thus, the difference observed between mature mRNA lengths of MISOG (3,341 bp) and SISOG (2,688 bp) (Table S1) can be attributed to the larger number of MISOG exons. On the other hand, MISOG introns (1,963 bp) were significantly ∼1.5 times longer than SISOG (1,336 bp) suggesting that the intronic regions, which are known to be longer than exonic regions in eukaryotes (7), are key MISOG/SISOG discriminative elements.

As previous studies on human and other eukaryotic genes have shown that the first intron is significantly longer than following ones (8–10), we performed an ordinal MISOG *versus* SISOG analysis for the 5’ and 3’ gene components (Fig. S1A, Table S7 and S8).

Both MISOG and SISOG exhibit a first 5’ intron (Fig. S1A; MISOG: 4,901 bp; SISOG: 2,953 bp) significantly longer than following introns (MISOG: 1,847-3,032 bp; SISOG: 1,341-1,823 bp). However, although introns lengths are divergent, MISOG displayed a much longer first 5’ intron than SISOG, especially concerning the upper quartile (MISOG: 19,992 bp; SISOG: 10,314 bp). We observed that the first MISOG and SISOG exons (Fig. S1B, Table S7) are longer (MISOG: 164 bp; SISOG: 195 bp) than following exons (MISOG: 123-125 bp; SISOG: 130-141 bp) and that the first MISOG and SISOG TERs (Fig. S1C, Table S3) are shorter (MISOG: 94 bp; SISOG: 117 bp) than the following TERs (MISOG: 119-124 bp; SISOG: 124-133 bp), suggesting that 5’ UTRs might account for the size discrepancies. Concerning the 5’ UTRs (Fig. S1D, Table S7), a decreasing length between first and second UTR exons is observed for both MISOG and SISOG. However, the first two 5’ UTR exons were found to have significantly longer lengths for MISOG (first and second exons, respectively 129 bp and 70 bp) compared to SISOG (first and second exons, respectively 97 bp and 39 bp). In addition, we noted a huge decrease in SISOG UTR’ numbers for positions 3-5 (241 /67 /26) compared to positions 1-2 (6,197 /1,546).

Concerning the last five 3’ components, no major differences between MISOG and SISOG were observed for introns regardless of the position (Fig. S1E, Table S8). Both MISOG and SISOG exhibit longer last exons (Fig. S1F; MISOG: 1,537 bp; SISOG: 1,221 bp) and last TERs (Fig. S1G; MISOG: 135 bp; SISOG: 158 bp) than the preceding ones (exons: MISOG: 122-124 bp; SISOG: 129-132 bp; TER: MISOG: 120-121 bp; SISOG: 125-130 bp). Finally, we noted that both MISOG and SISOG exhibit a single long 3’ UTR exon (Fig. S1H). However, 3’ UTR median exon length was 40% longer for MISOG (1,285 bp) compared to SISOG (918 bp).

To better understand the disparities between 5’ and 3’ non-coding regions, we analyzed the UTR in more detail and found that 75% of SISOG and only 41% of MISOG exhibit a single 5’ UTR exon attached to the CDS (Fig. 2A). Thus, most MISOG (59%) present multiple 5’ UTR exons, with the first one being the longest. At the opposite 3’ end, almost all MISOG and SISOG present a single long 3’ UTR exon attached to the CDS (Fig. 2B), confirming previous observations (10,11). These results are confirmed by considering the total UTR exon length for all genes (Fig. S2), where an 84% increase is observed for MISOG (MISOG: 204 bp; SISOG: 111 bp) at the 5’ UTR. Additionally, the length separating the TSS from the start codon was found to be 11 times longer for MISOG (1,577 bp) than for SISOG (143 bp) (Fig. 2C,Table S9).

**Figure 2.**
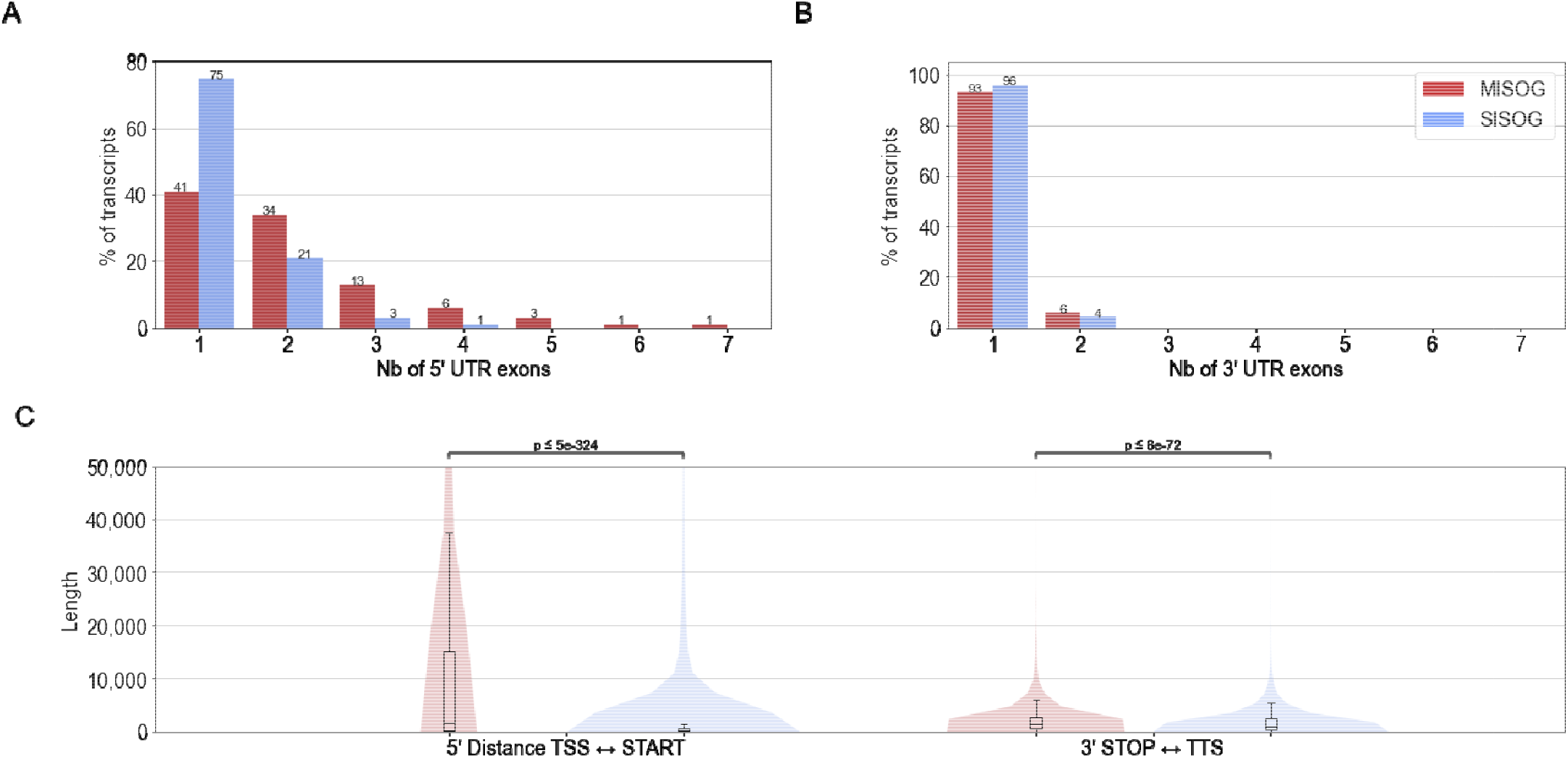
Comparative analysis of MISOG and SISOG 5’ and 3’ non-coding regions. (A-B) barplots indicate the percentage of MISOG/SISOG transcripts exhibiting from one up to seven 5’ or 3’ UTR exons. (C) Comparison of the MISOG *versus* SISOG median distances separating TSS from START codon (left) and STOP codon from TTS (right) (all values are available in Table S9). (A-B) represent the percentage of transcript according to the number of exons in 5’ (A) or 3’ UTR(B).

Finally, by analyzing the exon and TER frequencies across transcript isoforms in MISOG, we found that the closer these elements are to the 5’ end, the more likely they are alternatively spliced (Fig. S3 & Table S10). Indeed, the first alternative exon and the first alternative TER are present in only 50% of the isoforms. Moreover, second and third TER are also more alternatively spliced (resp 67% and 86%). Concerning the 3’ end, the last TER is found in 67% of isoforms and the last exon in 75%, corresponding to known 3’ UTR modulation (10).

## Conclusions

In this study, we have shown that 64% of HPCG in the RefSeq database are MISOG, representing 74% of the total genome length covered by coding genes (∼1,25 Gbp) and despite a large distribution, the detailed statistical analysis of the number and length of MISOG *versus* SISOG components (exons, TER, UTR and introns) revealed discriminative profiles (Fig. 1). Unsurprisingly, SISOG (n=7) have fewer exons than MISOG (n=12), but all SISOG exons are totally or partially coding while MISOG present at least one 5’ UTR exon without any coding part. Thus, the first 5’ SISOG and MISOG introns are clearly distinct, since in SISOG it is embedded in the coding region while in MISOG it is flanked by UTR. Such a distinct flanking environment may echo our observation that 5’ MISOG exons are statistically more prone to alternative splicing than the following exons. Finally, we show that the length of the first MISOG intron is almost 2 times longer than the SISOG one. The complex 5’ UTR architecture combined with a longer 5’ intron induces a fundamental difference of the TSS to start codon distance, which is generally 11 times longer in MISOG than in SISOG.

Various statistical studies have described the specificity of the 5’ eukaryotic gene region, notably concerning the properties of the first introns in term of length (8,9), conservation (11) or UTR inclusion (12,13) as well as the presence of *cis-*regulatory elements (14–16) and varying promoter number (17,18). For instance, it has already been noted that intron length impacts transcriptional time delay (19–21) or that the presence of 5′-UTR introns influences the choice of the alternative mRNA export pathway and the final mRNA targeting to endoplasmic reticulum and mitochondria (12). Thus, reanalyzing these distinctive properties in the light of the MISOG/SISOG profiles may allow a better discrimination of the processes and evolution linked to the splicing from those associated to alternative splicing.

Finally, this survey took advantage of the extended and high-quality of human data, and it remains to be seen whether applying the same protocol to other eukaryotic organisms with comprehensive isoform information may corroborate our results. Nevertheless, the finding of discriminatory features is promising as it opens the way to new *in silico* approaches to better predict the transcript isoform gene status and improve genome annotation.

All HPCG data are free to access and structured according to MISOG/SISOG status and all programing notebooks are provided and reusable (see Availability of data and materials).

## Methods

All definitions and methods described in the following sections are illustrated in Fig. S4.

### Data loading and processing

The RefSeq database (5) was selected as source of the genomic data for its associated quality (good annotation, non-redundancy, and low number of errors). The Human GFF file based on GRCh38p.13 assembly (version 109.20210514) was downloaded and filtered to retain only protein-coding genes and their features.

Using a python (3.7) script (*script: prepare_refseq*.*py*) based on the pandas library, four intermediate files were created to separately capture the four types of biological concepts (Fig. S4A) associated with protein-coding genes from the GFF files: genes (*feature=gene*), mRNAs (*feature=mRNA*), exons (*feature=exon*), and Translated Exonic Regions (TER) (*feature=CDS*).

Three main filtering steps were applied to curate the data. First, only entries having an identifier prefix of *NC*_ were retained. Second, only protein-coding genes were selected (*gene_biotype=protein_coding*). Third, for mRNA, exons and TER features, only curated entries with accession prefixes or parent entities starting with *NM*_ or *NP_* were retained.

Of the 42,793 gene entries (file G), 19,456 genes had a prefix NC_ and the attribute *gene_biotype=protein_coding*. After filtering 171 genes with no validated mRNAs (prefix NM_), the total dataset included 19,285 genes. We also excluded 1,983 single-TER genes, comprising 1,610 SISOG (of which 995 were single-exon genes) and 373 MISOG where the AS occurs uniquely in the UTRs.

Gene, mRNA, exon and TER lengths were calculated using the retrieved genomic coordinates. Gene start and end coordinates correspond to Transcription Start (TSS) and Transcription Termination (TTS) Sites respectively.

### Definition of MISOG and SISOG

We identified MISOG and SISOG based on the number of mRNAs per gene (respecting previously defined filtering steps). MISOG have at least two non-identical mRNAs while SISOG have only a single mRNA as presented in Fig S4B.

### Characterization of gene elements: exons, TER, introns and UTRs

Exons and TER lengths were computed using their respective start and end genomic coordinates. Intron boundaries were computed from exon positions as presented in the Fig. S4C. Based on previous observation (7), introns shorter than 26 bp were not analyzed.

Identification of 5’ /3’ UTR exons was determined using the transcription strand (+ = forward; - = reverse). UTRs were retrieved for each transcript by comparing exons and TER using a nested loop join algorithm conditioned by the presence of an overlap between an exon and a TER element. If so, UTR boundaries (excluding TER part of the exon) were computed as presented in the Fig. S4D. UTR boundaries were used as a new column in the file listing exons and TER for each mRNA.

For MISOG, gene elements may appear multiple times in RefSeq GFF file due to the data format structuration. To produce robust statistics, we identified and did not consider redundant elements (see Fig. S4E).

### Gene element ordinal position

For each gene element, the ordinal position was defined as presented in Fig. S4F with respect to the transcription strand (+ = forward; - = reverse) and its position in the mRNA.

### Exons and TER usage frequencies across isoforms

For MISOG, we identified and counted the constitutive (present in all mRNAs) exons and TERs and the alternative ones (present in a subset of mRNAs). Frequencies were computed as presented in Fig. S4G and defined as the number of transcripts where the exon/TER is present divided by the total number of transcripts in the gene.

### Statistical analyses

As parametric statistics (mean and standard deviation) are highly sensitive for extreme values and due to the widespread distribution of gene element lengths across all HPCG, all comparisons were performed using non-parametric statistics (median and quartiles). Statistically significant differences were evaluated using standard Mann-Whitney U tests.

## Abbreviations

CDS: Coding DNA Sequence
TSS: Transcription Start Site
TTS: Transcription Termination Site
UTR: UnTranslated Region
HPCG: Human Protein Coding Genes
MISOG: Multiple transcript ISOform Genes
SISOG: Single transcript ISOform Genes
NMD: Non-sense Mediated Decay

## Declarations

### Ethics approval and consent to participate

Not applicable.

### Consent for publication

Not applicable.

### Availability of data and materials

The code used in this study can be found at https://github.com/weber8thomas/MISOG_SISOG under MIT License. RefSeq raw GFF file was retrieved from FTP site (GCF_000001405.39). The data sets are accessible in compressed CSV, Apache parquet and XLSX formats at https://zenodo.org/record/5546587. All developed programs and notebook were specifically designed to allow facilitated updates on new versions of RefSeq GFF files.

### Authors’ contributions

T.W., L.M, K.C., and O.P. designed the study. K.C. and O.P. supervised the work. T.W. and N.S. produced the visualizations. T.W., K.C. wrote the manuscript. J.T. and O.P. contributed to revision of the manuscript.

## Acknowledgements

No statement

