## Supplementary materials for "Do 5’ regions of human protein-coding genes contain the blueprints for alternative splicing?"

**Supplementary Material**

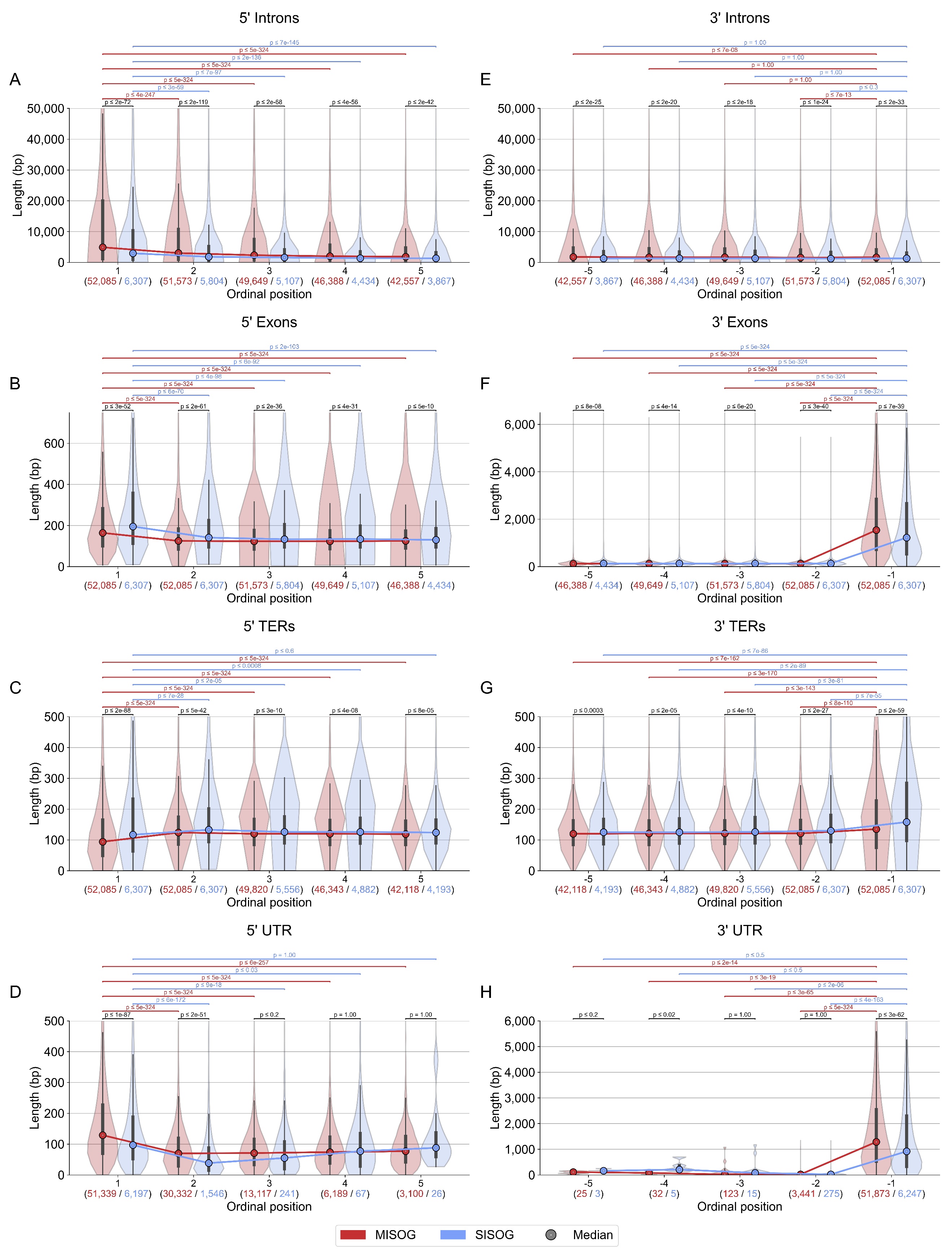

**Figure S1 – Length distribution of gene elements (Intron, Exon, TER and UTR exons) according to ordinal positions.**

All plots represent the length distribution of gene elements according to ordinal position (as illustrated in Fig. S4F). Plots A-D correspond to the first five 5’ elements in the genes (1 up to 5), plots E-H correspond to the last five 3’ elements (-5 to -1). The p-values obtained through Mann-Whitney U tests are displayed above the violin plots. Black p-values correspond to comparisons between MISOG and SISOG while colored p-values represent comparisons between first elements and following ones (MISOG: red, SISOG: blue). Values under each plot represent the numbers of observed elements at the ordinal position for MISOG (red) and SISOG (blue).

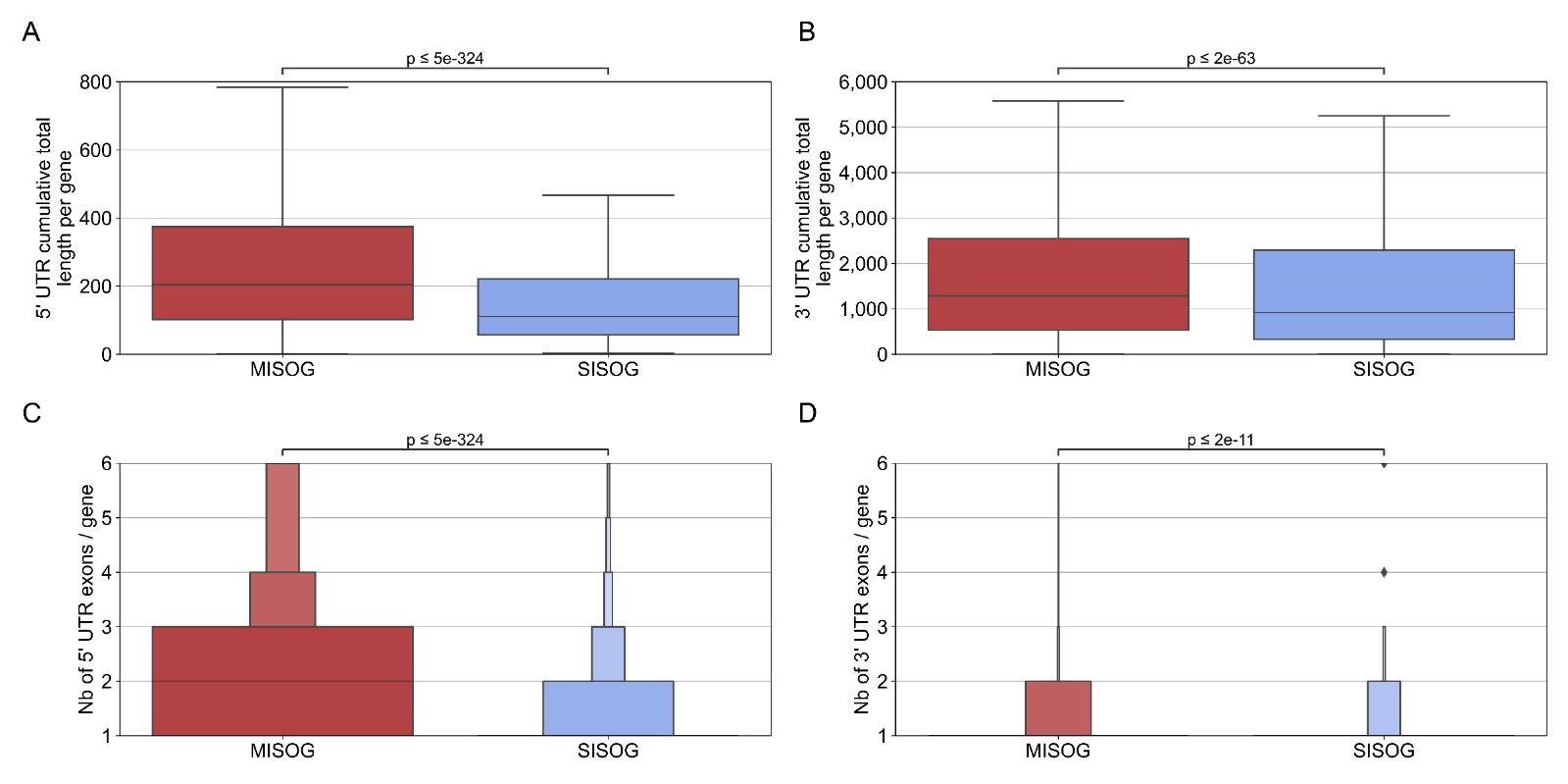
**Figure S2 – Cumulative length and number of 5’ and 3’ UTR exons by gene**

Cumulative total lengths of 5’ (A) and 3’ (B) UTR regions per gene. Number of 5’ and 3’ UTR exons per gene are displayed in (C, D). The p-values obtained through Mann-Whitney U tests are displayed above the boxplots (A, B) and boxenplots (C, D).

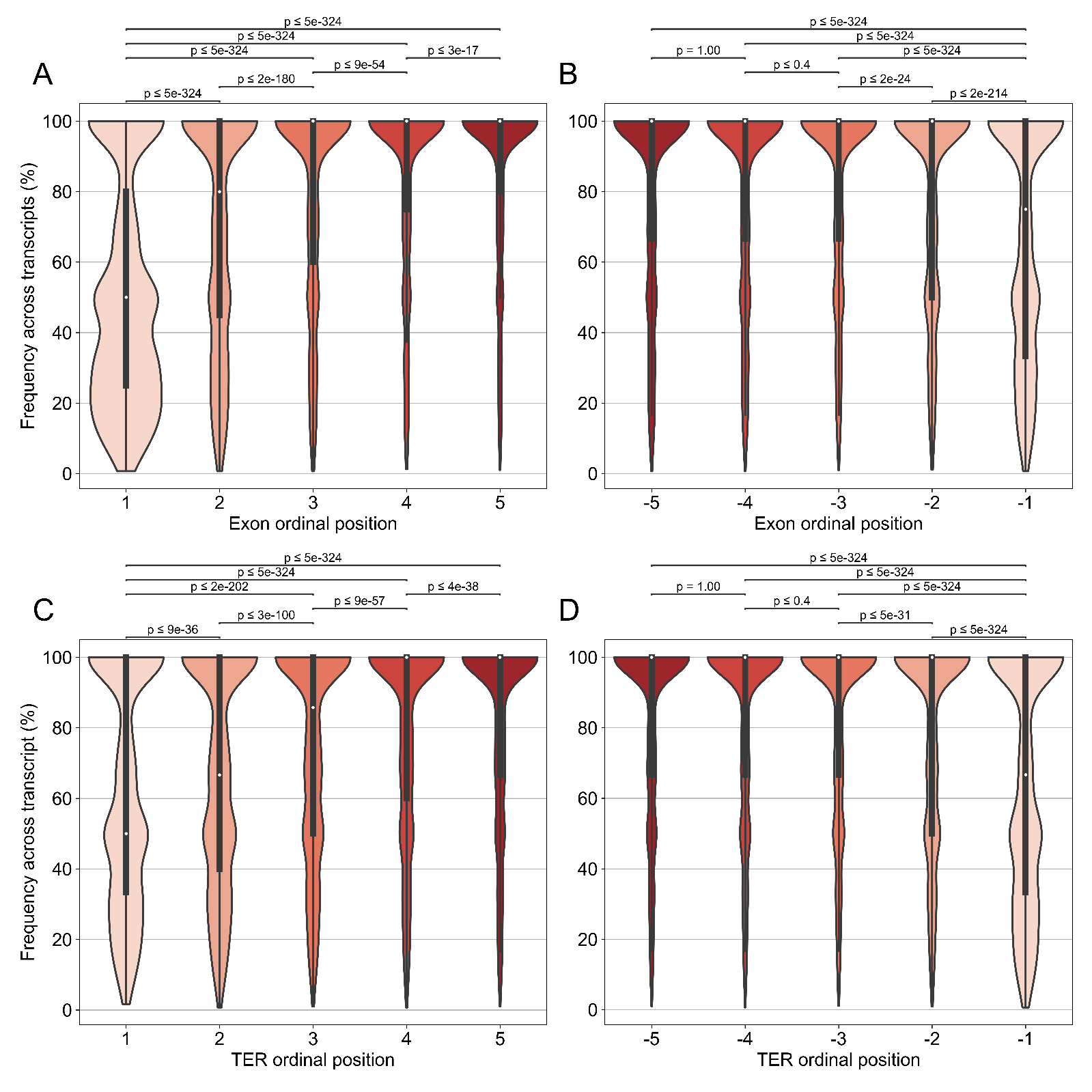

**Figure S3 – Percentage of alternative exons and TERs according to their ordinal positions.**

All plots show calculated frequencies of alternative exons (A, B) and alternative TERs (C, D) as illustrated in Fig. S4G. Plots (A, C) correspond to the first five 5’ elements in the genes (1 up to 5), while plots (B,D) correspond to the last five 3’ elements (-5 to -1). The p-values obtained through Mann-Whitney U tests are displayed above the violin plots.

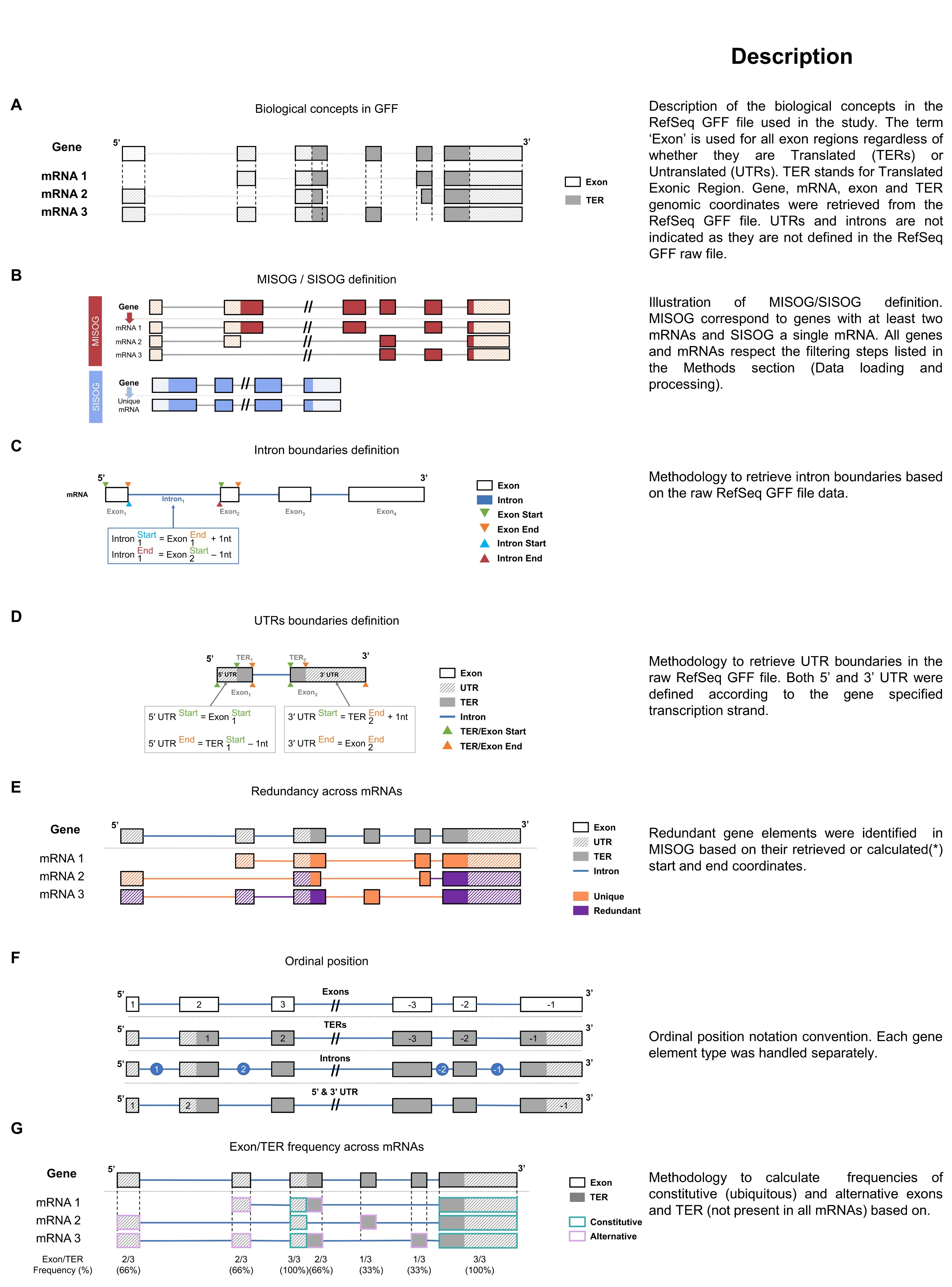

**Figure S4 – Material and Methods schemas**
